# Coevolutionary Stability of Host-Symbiont Systems with Mixed-mode Transmission

**DOI:** 10.1101/2023.02.06.527336

**Authors:** Nandakishor Krishnan, Lajos Rózsa, András Szilágyi, József Garay

## Abstract

The coevolution of hosts and symbionts based on virulence and mode of transmission is a complex and diverse biological phenomenon. We introduce a conceptual model to study the stable coexistence of an obligate symbiont (mutualist or parasite) with mixed-mode transmission and its host. The existence of evolutionarily and ecologically stable coexistence is analyzed in the framework of coevolutionary dynamics. Using an age-structured Leslie model for the host, we demonstrate how the obligate symbiont can modify the host’s life history parameters (survival and fecundity) and the long-term growth rate of the infected lineage. The evolutionary success of the symbionts is given by the long-term growth rate of the infected population (multi-level selection). When the symbiont is vertically transmitted, we find that the host and its symbiont can maximize the long-term growth rate of the infected lineage. Moreover, we provide conditions for the ecological and evolutionary stability of the resident host-symbiont pair in the coevolutionary model, which does not allow invasion by any rare mutants (each mutant dies out by ecological selection). We observed that ecological competition, clearing of infection, and density-dependent interactions could play a role in determining the criteria for evolutionary stability.

## 1. Introduction

The coevolution of hosts and symbionts (mutualists and parasites, including pathogens) constitutes a complex and diverse biological phenomenon (Clayton et al. 2015; Gandon et al. 2008). The symbionts’ virulence and modes of transmission are two factors that significantly influence these processes. Virulence is usually defined as the parasites’ ability to reduce infected hosts’ survival and reproductive success. In our interpretation, mutualists exhibit negative virulence since they increase (rather than decrease) infected hosts’ survival and reproductive success. The modes of transmission of symbionts to the following host individual are typified according to the genetic relatedness between the former and the next host. Transmission from parent to offspring (in a more general form, transmission between close genetic kins) is called vertical transmission. Contrarily, we define horizontal transmission as the transfer of symbionts between genetically non-kin hosts. Most symbionts use a combination of vertical and horizontal transmission, which we call a mixed-mode transmission. A massive body of theoretical and empirical evidence unequivocally suggests that symbionts’ virulence and mode of transmission are interrelated characteristics (Ebert 2013; Ewald 1987). Exclusive vertical transmission is frequently found only in mutualistic interactions (Bright and Bulgheresi 2010). In this case, infected host populations outcompete non-infected ones; thus, the infection reaches fixation in host populations (Ewald 1987). However, imperfect vertical transmission (when some offsprings fail to inherit the symbiont) can prevent symbiont fixation (Afkhami and Rudgers 2008). Contrarily, symbionts that transmit exclusively (or primarily) horizontally (like vector-borne and water-borne pathogens, mobile parasites, etc.) tend to be highly virulent, even lethal (Ebert 2013; Ewald 1987). The exclusively horizontal or exclusively vertical transmission systems are just the extremes of a continuum, while most of the real-life host-symbiont systems are characterized by mixed transmission. Hereafter we consider only such mixed-mode transmissions in host-symbiont systems. The symbionts’ effect on the host’s life history may also be quite complex; infections may increase or decrease host mortality, fecundity, or both. These effects may markedly differ between different developmental stages of the host; for instance, in the case of COVID-19, mortality differs between young and adult individuals (Liu et al. 2020).

Our paper focuses on the evolutionarily stable coexistence of mixed (vertical and horizontal) transmission symbionts (such as parasites, pathogens, and mutualists) and their host. We only consider obligate (with no free-living stages) and host-specific (connected to a single host species) symbionts. Furthermore, we consider the aging of the hosts, i.e., hosts have different developmental stages (juvenile and adult stages) and a finite lifespan. We investigate how symbionts of different types can modify a host’s life history. The conceptual model introduced in the coming sections focuses on the host-symbiont interactions and the symbionts’ effect on the survival rate and the fecundity of the infected host population. To ensure the ecological coexistence of infected and non-infected host populations, we consider density-dependent interactions, ecological competition, and horizontal infection between the populations. Moreover, we must also consider that infected hosts can lose their symbionts in a density-independent way, e.g., by stochastic loss or recovery from infection – often called ‘clearing.’ In our view, three mechanisms play a significant role in the coevolution of the hosts and their symbionts: 1) the nature of the host-symbiont interaction (how infection affects host mortality and fecundity), 2) the proportion of vertical and horizontal transmission, and 3) clearing of infected hosts. These mechanisms may not evolve independently. For example, in the case of mutualism, vertical transmission and low clearing rates (often zero) benefit both species. Contrarily, in the case of parasitism, the hosts benefit from getting rid of their symbionts (recovering from infection) or avoiding infections (both horizontal and vertical). To focus exclusively on the demographics of the infected population as defined by the host-symbiont interactions, we apply the following *uniformity conditions:* 1) the transmission mode (proportion of vertical versus horizontal routes) is independent of the host phenotype, 2) the frequency of clearing is also independent of the host phenotype, and 3) there is no difference in the competitive ability between different populations.

We have two objectives that we would like to attain in this paper. Firstly, we are looking for an evolutionary stable host-symbiont pair in the framework of a coevolutionary ecological model (Cressman and Garay 2003a, b) which does not allow any rare mutant to invade the resident system (Maynard Smith and Price 1973). From a mathematical point of view, we give the evolutionary stability condition by the local asymptotical stability of the coevolutionary differential equations combined with the uninvadability of rare mutants into the stable resident system. Secondly, we are interested in the possible connection between the Evolutionarily Stable Strategy (ESS) definition in game theory (Maynard Smith and Price 1973) and the stability conditions of our coevolutionary dynamics. Here we focus on the strict Nash equilibrium game solution, which claims that no mutant can increase its benefit at the ESS by unilaterally changing its strategy.

The remaining part of the paper is organized into several sections. Section 2 introduces the preliminaries for the model; section 3 describes the effect of infection on the life history of the host; section 4 deals with the description and stability analysis of the resident system; section 5 introduces mutants into the resident system; section 6 analyses the possible connection between our dynamical model and game theory, and finally section 7 concludes the manuscript with discussions, inferences, and biological examples.

## 2. Methods

The general framework of our selection situation belongs to the *multi-species group (multilevel) selection* models (Simon 2014) since the host, together with its symbionts, form a well-defined ‘group’ (namely an infected individual). This group selection model has the following main features:

1. *Group formation process:* groups can reproduce by means of host reproduction and vertical transmission. Horizontal transmission may change non-infected host lineages to infected ones; thus, new groups can appear. Finally, groups may dissolve due to clearing. The novelty of this paper is the inclusion of mixed-mode transmission and clearing. We assume that the group formation dynamics is based on host-symbiont interactions. The horizontal infection and clearing are independent of the survival rate and fecundity of the infected populations. Note that the dynamics of the model is connected to the standard epidemic models (Hethcote 2000; Kermack and McKendrick 1927; Tsay et al. 2020), focusing on horizontal transmission, which only connects infected and non-infected individuals. The main reason we cannot simply apply the standard epidemic models is the inclusion of vertical transmission, which connects the generations.
2. *Group payoff:* the symbionts’ and hosts’ fitnesses together determine the infected population’s evolutionary success. Moreover, the evolutionary success (fitness) of the symbionts – since they are obligate – is given by the evolutionary success of the infected population. Thus, ‘groups’ have a well-defined payoff, i.e., the long-term growth rate of the infected population.

Since the obligate symbiont and its host together determine the evolutionary success of the infected host population, the evolutionary success of vertically transmitted parasites or mutualists can be obtained by following the infected host lineages. Game theory is thus one of the inevitable methods that can be applied to the model (Ezoe 2009; Noë and Hammerstein 1995). However, this ‘shared interest’ (similar to group selection theory) makes the direct application of the evolutionary matrix game challenging (Ezoe 2009; Maynard Smith and Price 1973). The adaptive dynamics method (Dercole and Rinaldi 2008; Genkai-Kato and Yamamura 1999) can also be applied to this problem but cannot be generalized if the number of species and age-classes are high. We now mention here a few relevant models on the evolutionary stability of vertically transmitted symbionts (Akçay 2015; Friesen and Jones 2012) without providing a comprehensive literature overview. Ferdy and Godelle 2005 investigated how symbionts’ transmission mode (horizontal or vertical) and virulence should coevolve, and the basic structure of their ecological model is similar to ours. Some studies investigated how host demography affects host-symbiont coexistence using real-life examples (Afkhami and Rudgers 2008; Bibian et al. 2016; Chung et al. 2015). We consider that the infected host’s life-history parameters (such as survival rate and fecundity) at a time are determined by the symbiont and the host. Our starting point is the well-known Leslie demography model (Bibian et al. 2016; Caswell 2001; Charlesworth 1980); thus, the long-term growth rates of the different populations are given implicitly. We utilize our formerly published *Kin Demographic Selection Model* (Garay et al. 2016) for a single species. The three main assumptions of this model are: 1) different phenotypes are determined by their hereditary life-history strategies, 2) the survival rate depending on total density is the same for all individuals, independent of their phenotype and age, and 3) the behavior within the kin and the density-dependent survival process is independent. The main idea behind this model is that the evolutionarily best phenotype maximizes its phenotypic long-term growth rate. We assume age-structured Leslie models where phenotype A and B can be described by their population vectors (containing the number of individuals at *n* different ages): 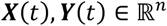, respectively, and populations of A and B are governed by the corresponding Leslie matrices: 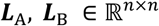. The discrete dynamics for phenotype A is: ***X***(*t* + 1) = ***L***_A_***X***(*t*), and similarly, for phenotype *B*. Population vector ***X***(*t*) asymptotically tends to the equilibrium age-structure distribution represented by the leading eigenvalue of the Leslie matrix. The corresponding leading eigenvalue *λ*_A_ (which is the dominant eigenvalue of ***L***_A_) defines the long-term growth rate of the population. If *λ*_A_, *λ*_B_ > 1, both populations grow and if *λ_A_* > *λ_B_*, the relative frequency of phenotype B tends to zero. Indeed, according to the original Darwinian view, we need some degree of density-dependent selection to keep the total density of these two phenotypes at the carrying capacity *K* (Garay et al. 2016). Now, by selection, the total density of the system reduces to *K* proportionally, i.e.,

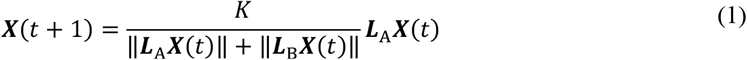

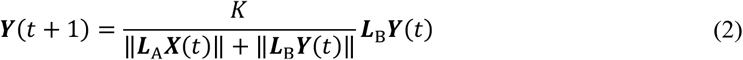

where, ∥***X***(*t*)∥ = ∑*_i_ X_i_*(*t*), denotes the sum of elements of the vector.

Since our main focus is on how the symbionts modify their hosts’ survival rate and fecundity, we apply the *uniformity conditions;* namely, the density-dependent competition, the horizontal transmission, and the clearing are all independent of the host’s age to the new model. In addition, while we considered phenotypes of the same species in the former model, here we generalize it for the interactions between an obligate symbiont and its host, where symbionts can modify their hosts’ life history. Namely, parasites and pathogens decrease their hosts’ fecundity and/or survival rate, while mutualists increase that. Clearly, we face a coevolutionary problem since the obligate host-symbiont system has two interacting ‘species,’ represented here as infected and non-infected host populations. Since we are looking for an evolutionarily stable ecological state, following Maynard Smith and Price 1973, we also assume that the mutations (heritable changes in host strategies or characters) are rare enough. There are two consequences of this assumption. First, after a mutation appears in the system, there is enough time for natural selection to eliminate the least fit types. Second, we need to consider only the case when there is only a single mutant type in each population because the population reaches its stable rest point before the next mutation arises. The original verbal definition of the evolutionarily stable strategy (ESS) with a frequency-dependent interaction scheme is the following for a single species. If the overwhelming part of the population uses this strategy (or phenotype), then a rare mutant cannot invade the population (Maynard Smith and Price 1973). However, in the present study, we consider two species (host and symbiont) with ecological competition between infected and non-infected host populations; thus, the original one-species ESS definition cannot be directly applied. To create an ESS concept suitable for our present study, we apply the *N*-species evolutionary stability concept (Cressman et al. 2020; Cressman and Garay 2003a, b; Garay 2007). This concept belongs to the general theory of coevolution (that combines effects of ecology and evolution) based on arbitrary nonlinear fitness functions and a finite number of phenotypes. The primary view of this concept is the following: since mutations are rare enough, there is enough time for the resident system to reach the stable state by ecological selection, in other words, by ecological dynamics. Thus, each mutation appears in the ecologically stable state of the resident system. In other words, the time scale of mutation is much longer than the ecological time scale. Furthermore, mutations increase the dimension of the system since new types appear. Now the question arises whether any mutant can invade the resident system at its ecological rest point. Following the primary idea by Maynard Smith and Price 1973, we consider a resident ecological system to be evolutionary stable if rare mutants cannot invade the system according to ecological dynamics. In the next section, we utilize the Leslie model (Metcalf and Pavard 2007) and introduce the strategy-dependent life history model.

## 3. Strategy dependent discrete life history model

The two populations considered in the Kin Demographic Selection Model (Garay et al. 2016) were independent, while in the present model, the infected and non-infected populations are not independent since the infected individuals can be cleared (getting rid of symbionts), and non-infected individuals may get infected through horizontal transmission. We consider that the infected host’s life-history parameters (survival rate and fecundity) at a time are determined by the symbiont and the host together. Thus, we first generalize the former model to consider the connections between the infected and the non-infected populations.

### 3.1 Life history model of the non-infected host

For simplicity, we assume that the host’s lifespan is only three years. In the first year, juvenile hosts do not reproduce. *α*_1_, *α*_2_ > 1 denote the average fecundity of the host during the second and third years, respectively. Note that the Leslie model only follows individuals (females) that can produce offsprings. Further, 0 < *ω*_0_ < 1 and 0 < *ω*_1_, < 1 denote the average survival rate from juvenile to one-year-old host and from one-year-old to two-year-old host, respectively. The 3-year-old hosts die after reproduction. Using these notations, the standard Leslie model for a non-infected host population is:

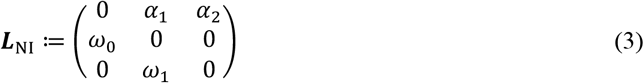

The long-term growth rate of the non-infected population is given by the leading eigenvalue (*λ*_NI_) which is the unique, dominant, positive, and real solution of the following characteristic polynomial of the matrix ***L***_NI_,

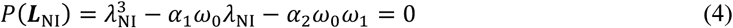

In our model, we consider a trade-off between host fecundities and survival rates (Garay et al. 2016). Let *s* ∈ [0,1] denote the host’s resource allocation strategy. Introducing strategy-dependent fecundity functions *α*_1_(*s*), *α*_2_(*s*) ≫ 1, which decreases with increasing *s* and strategy-dependent survival rates *ω*_0_(*s*), *ω*_1_(*s*): [0,1] → (0,1), which increases with increasing *s*, we get the following strategy-dependent Leslie matrix for the non-infected population,

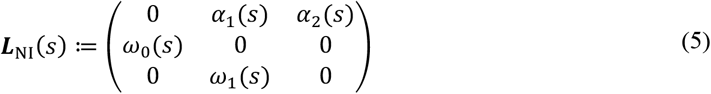

#### Example 1

Consider the simplest trade-off in the form of linear functions *α*_i_(*s*) = *α*_1_ – 75*s*, for *i* = 1,2, *ω*_0_(*s*) = *ω*_0_ + 0.75*s* and *ω*_1_(*s*) = *ω*_1_ + 0.65*s*, we have:

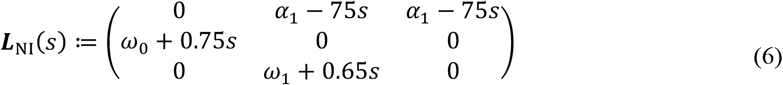

For simplicity, throughout this paper we fix the weights of this trade-off (i.e., 0.75, 0.65 and 75), where 0 < *ω*_0_ + 0.75*s* < 1 and 0 < *ω*_1_ + 0.65*s* < 1 for *s* ∈ [0,1]. Now we look for 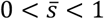 which maximizes *λ*_NI_(*s*) (see Figure 1), when *α*_1_ = 100, *ω*_0_ = 0.2, *ω*_1_ = 0.3. The corresponding characteristic polynomial is,

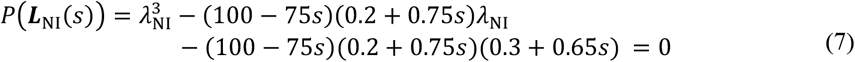

**Fig. 1.**
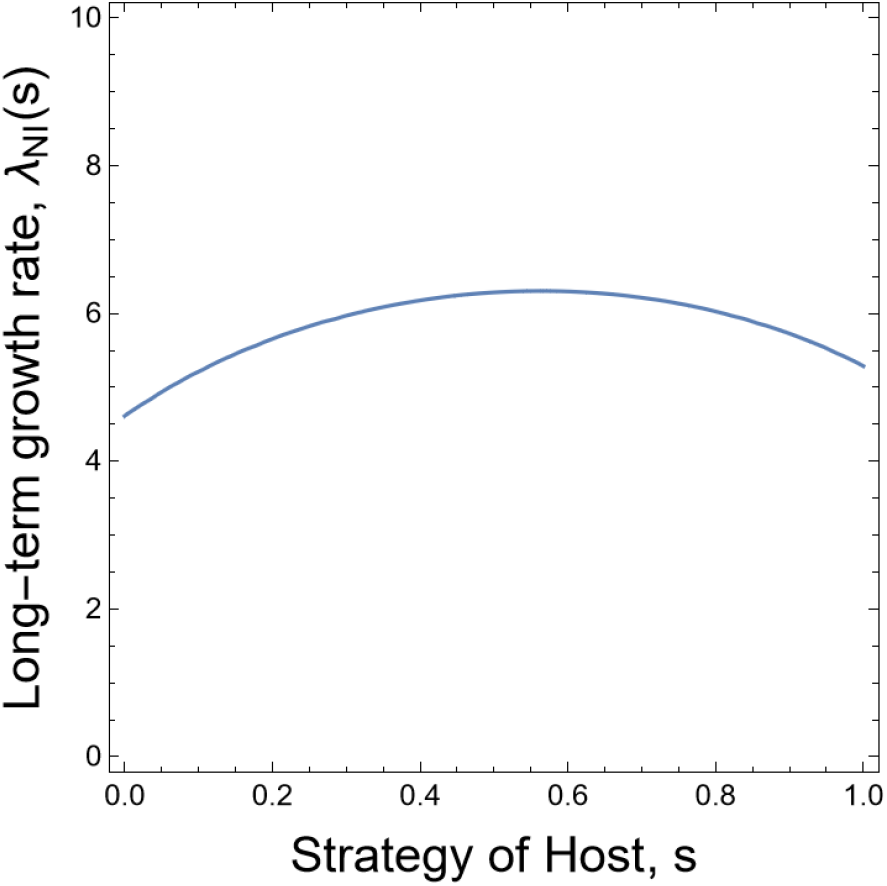
Optimization of the long-term growth rate of the non-infected host population (*λ*_NI_(*s*)) with respect to the host strategy (*s*). The maximum value is at 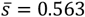

### 3.2 Life history model of the infected host

Now we consider the symbiont-infected host population. The symbionts can modify the hosts’ fecundity and survival. This modification strategy is denoted by *σ* ∈ [0,1]. For a fixed *σ*, the symbionts’ effect on the fecundity of their host is given by a function *f_i_*(*σ*), where the index denotes different cohort (age class or developmental stage). Similarly, the function *g_i_*(*σ*) gives the symbionts’ effect on their host’s survival rate. In the case when hosts have no alternative strategies (for fixed *s*), we get the following general model for the infected host lineage with vertically transmitted symbionts,

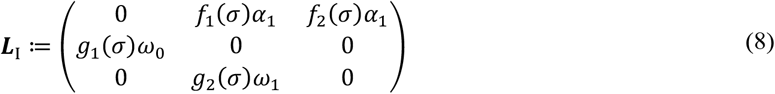

Note that a trade-off can be considered in this model when the functions *f* and *g* increase or decrease with respect to *σ* ∈ [0,1].

#### Special case

Consider the case when the functions *f* and *g* are constants, i.e., let 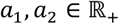 and 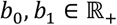 such that *f*_1_(*σ*) = *a*_1_, *f*_2_(*σ*) = *a*_2_, *g*_1_(*σ*) = *b*_0_ and *g*_2_(*σ*) = *b*_1_ for each *σ* ∈ [0,1]. We get the following life-history model for the infected hosts,

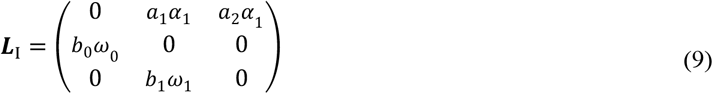

For the consistency of this model, we need 0 < *b_j_ω_j_* < 1 for *j* = 0,1, since *b_j_ω_j_* is the survival rate of *j*-th infected host age class. The following are the possible biological interpretations of positive parameters *a_i_* and *b_j_*:

1. If the symbiont is purely parasitic, i.e., decreases both survival and reproduction of its host species, we have 0 < *a_i_* < 1, 0 < *b_j_* < 1.
2. If the symbiont is purely mutualistic, i.e., increases both host survival and fecundity, we have *a_i_* > 1 and *ω_j_* < *bω_j_* < 1.
3. Mixed case, i.e., when the infection increases host survival but reduces its fecundity or vice-versa.

For simplicity, we assume that the vertical transmission is perfect. Let us now focus on the Leslie matrix ***L***_I_ with fixed parameters. Here the long-term growth rate of the infected host population (*λ*_I_) is given as the unique, dominant positive solution of the following characteristic polynomial of ***L***_I_,

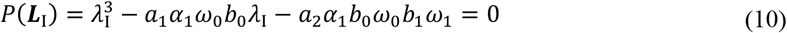

After simple rearrangement, we obtain the following equation for the eigenvalues,

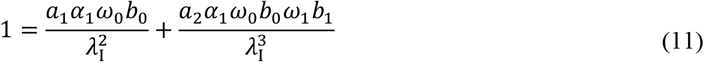

Eq. 11 shows that the long-term growth rate is a strictly increasing function of parameters *a*_1_, *a*_2_, *b*_0_, *b*_1_. An obvious consequence of these monotonicities is that the symbionts can increase their own evolutionary success by increasing their host’s fecundity and survival parameters.

#### Example 2

To demonstrate the possible effects of a symbiont species on its host species, we consider the following numerical example for the infected host population. Let us assume that the non-infected population has the following demographic parameters: *α*_1_ = *α*_2_ = 100, *ω*_0_ = 0.2, and *ω*_1_ = 0.3. The corresponding long-term growth rate of the non-infected population (i.e., the dominant eigenvalue of the Leslie matrix in Eq. 3) is *λ*_NI_ = 4.615. Consider the following Leslie matrix for the infected lineage,

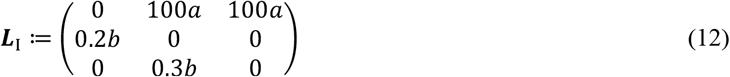

where, for simplicity, we assume that the effect on fecundities and survival rates are age-class independent, i.e., *a*_1_ = *a*_2_ = *a* and *b*_0_ = *b*_1_ = *b*. Figure 2 shows the curve of *b*(*a*) function based on Eq. 11 that satisfies *λ*_I_ = *λ*_NI_. Over this curve, where *λ*_I_ > *λ*_NI_, the infected population dominates the non-infected population.

**Fig. 2.**
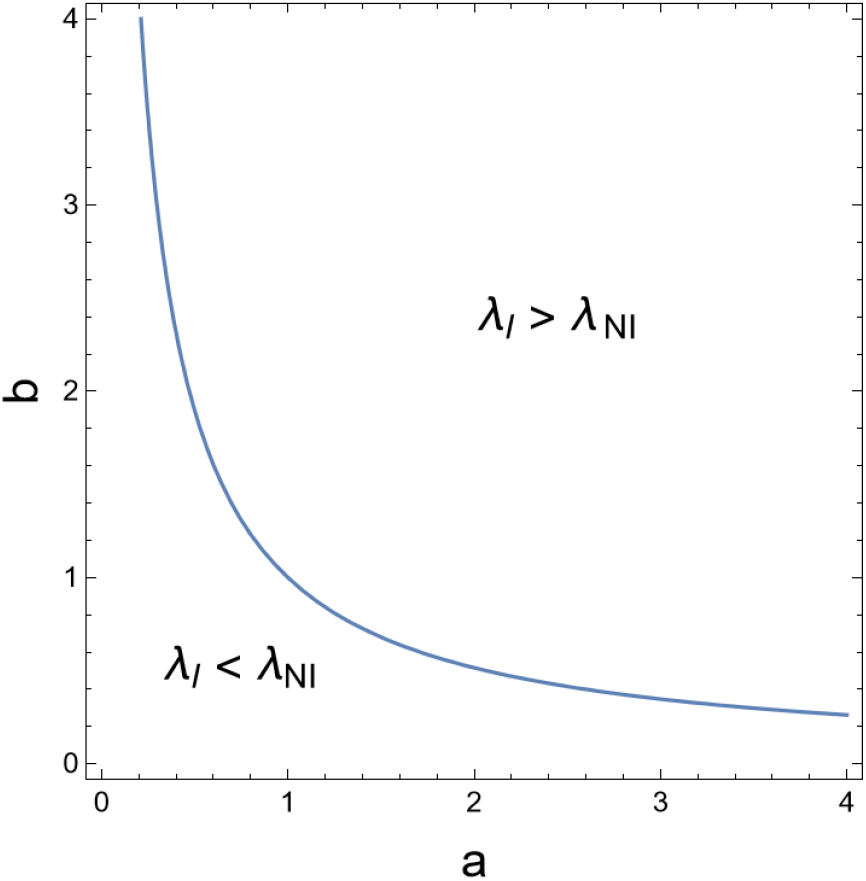
The curve denotes the parameter pairs *a* and *b* when the long-term growth rates of the two populations are the same (*λ*_I_ = *λ*_NI_). Over the curve, the long-term growth rate of the infected lineage is higher (*λ*_I_ > *λ*_NI_). Parameters are *α*_1_ = 100, *ω*_0_ = 0.2, *ω*_1_ = 0.3, and we assumed age-class independent effect on fecundities and survival rates

Now we give some numerical examples to demonstrate the symbiont’s effect on the long-term growth rate of the infected population. Assume *α*_1_ = *α*_2_ = 100, *ω*_0_ = 0.2, and *ω*_1_ = 0.3; thus the long-term growth rate of the non-infected population is *λ*_NI_ = 4.615.

1. Purely mutualistic: the symbionts increase the fecundity and survival rates of the hosts. For instance, if *a* = 2.5, *b* = 1.5, then *λ*_I_ = 8.877.
2. Purely parasitic: the symbionts decrease the fecundity and survival rates of the hosts. For instance, if *a* = 0.5, *b* = 0.5, then *λ*_I_ = 2.308. Our former model (Garay et al. 2016) predicted the extinction of parasitic symbionts because of the lack of horizontal infections.
3. Mixed strategy: the symbionts increase host survival but decreases host reproduction or vice-versa. This type of symbionts can either increase or decrease the long-term growth rate of the infected population. For instance, when the symbiont increases host reproduction but decreases host survival, i.e., *a* = 2.5, *b* = 0.5, *λ*_I_ = 5.073; the infected population has a higher growth rate (*λ_NI_* < *λ_I_*). Contrarily, when the symbiont decreases host reproduction but increases host survival, i.e., *a* = 0.5, *b* = 1.5, *λ*_I_ = 4.081; the growth rate of the non-infected population is higher.

### 3.3 Life history model of the infected host population with host and symbiont strategies

Let us now focus on the hosts’ and their symbionts’ common effect on the survival and fecundity of the vertically infected host population. As we have already considered, the host has a trade-off between fecundity and survival (*s* ∈ [0,1]) and the symbionts can modify infected hosts’ fecundity and survival rates (*σ* ∈ [0,1]). The generalized Leslie matrix corresponding to the model is,

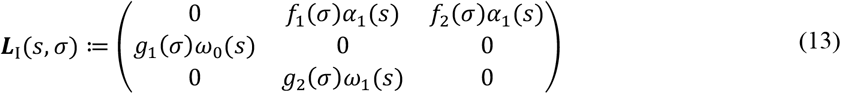

#### Example 3

Consider two trade-offs in the Leslie matrix of the above infected population,

1. Host trade-off between reproduction and survival (as in Example 1): let *α_i_*(*s*) = *α*_1_ – 75*s, ω*_0_(*s*) = *ω*_0_ + 0.75*s* and *ω*_1_(*s*) = *ω*_1_ + 0.65*s*.
2. Trade-off between the symbionts’ effects on host survival and fecundity: let *f*_1_(*σ*) = *f*_2_(*σ*) = 10*σ* and *g*_1_(*σ*) = *g*_2_(*σ*) = 1 – *σ*.

Now the Leslie matrix of the infected population is,

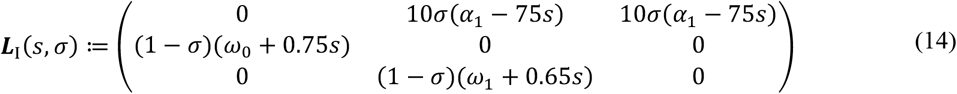

The long-term growth of the infected population and its optimization with respect to different values of *s* and *σ* are given in Figure 3.

**Fig. 3.**
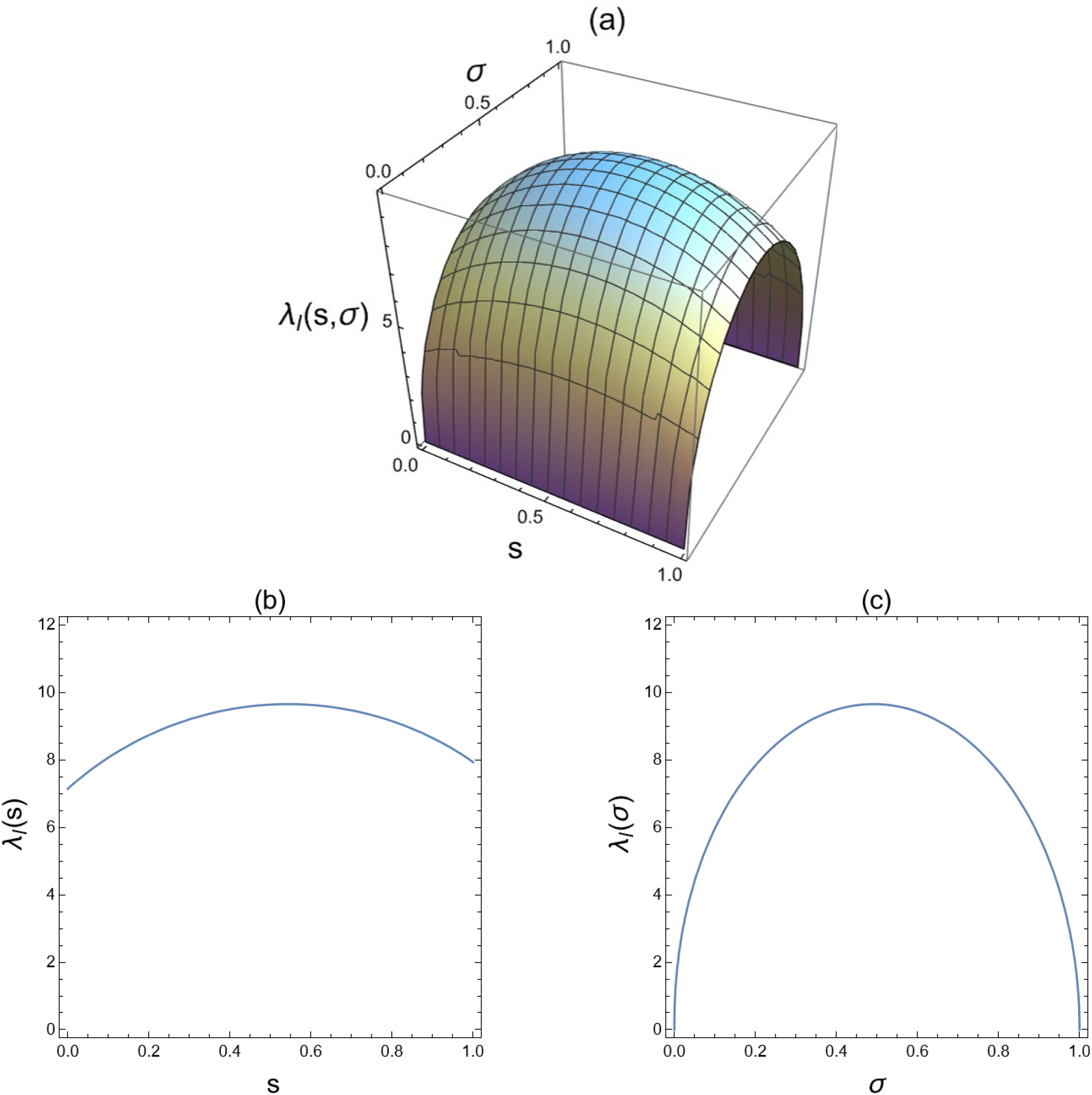
a) Long-term growth rate of the infected population on *s* – *σ* plane with maximum value 9.647 at (*s, σ*) = (0.544, 0.492); b) Maximization of the infected population’s long-term growth rate with respect to *s* for *σ* = 0.492; c) Maximization of the infected population’s long-term growth rate with respect to *σ* for *s* = 0.544

## 4. Competitive selection dynamics for the resident system

Based on the above discrete model of vertical transmission in infected and non-infected populations, we can derive the “ecological” dynamics. We introduce selection dynamics for the resident system with interacting infected and non-infected populations. Infected and non-infected populations interact in three ways: 1) by horizontal transmission (when a non-infected individual becomes infected (Ebert 2013), 2) by clearing (when an infected individual becomes non-infected), and 3) competition between individuals of the two populations. Implementing the interactions in the structured model results in high-dimensional systems with limited analytical tractability (Castelletti and Barbarossa 2020; Tian et al. 2018). To avoid these difficulties, we assume continuous-time population dynamics with the Malthusian growth rates derived from the structured model. The dynamics becomes two-dimensional by considering the dynamics of the two populations’ total densities. This simplification allows us to connect the dynamics of a structured population and the process of population regulation in a simple way, even though the high-dimensional model would yield more accurate predictions. To preserve analytical tractability, we set up a conceptual and qualitative model explaining the behaviour of the considered symbiotic systems without allowing quantitative validation. However, this makes it hard to use the standard invasibility plot method of adaptive dynamics theory (Dercole and Rinaldi 2008) when the matrix dimension is high. Therefore, we apply the coevolutionary ecological stability to find the evolutionarily stable resident system.

First, let us shed light on how we introduce the Malthusian growth rate into the dynamical system. In the discrete model, consider the non-infected and infected populations with *λ*_NI_ and *λ*_I_ respectively as long-term growth rates. Suppose the interconnection between these populations is uniform, i.e., density-dependent competition, horizontal infection, and clearing are independent of host’s age. We can use the well-known asymptotic behaviour of the Leslie model (Caswell, 2001), i.e., for each fixed Leslie matrix (independently of the initial state of the population), the population reaches an equilibrium vector (which is proportional to the eigenvector of the Leslie matrix) after enough time. In this case, the whole population’s growth rate is the Leslie matrix’s unique leading eigenvalue at its eigenvector (stable age distribution). The discrete dynamics of the non-infected population is governed by the following equation: ***x***_NI_(*t* + 1) = ***L***_NI_***x***_NI_(*t*), where ***x***_NI_(*t*) is the population vector of the non-infected population at time *t* and ***L***_NI_ is the Leslie matrix corresponding to the non-infected population. Based on the asymptotic behaviour of the Leslie model, the long-term growth rate of the non-infected population, denoted by *λ*_NI_ (the leading eigenvalue of ***L***_NI_) gives the population’s asymptotic growth rate. Similarly, we have the long-term growth rate of the infected population (*λ*_I_). We invoke the known fact that the logarithm of the growth rate of a Leslie model (Caswell, 2001) can be interpreted as the Malthusian parameter (*r*) of the corresponding continuous-time dynamics (*r* = ln *λ*). Based on this, we can consider ln(*λ*_NI_) and ln(*λ*_I_) as the Malthusian growth rates of the non-infected and infected populations under our uniformity conditions.

We denote 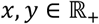 as the total population size of the non-infected and infected populations respectively, i.e., the total size of the population near to the stable asymptotic state of the respective Leslie matrices. Further, we denote the horizontal transmission rate by *β* ∈ (0,1). For simplicity, we assume that the horizontal transmission rate does not depend on the individual’s age. Furthermore, at the state when there is *x* non-infected and *y* infected individuals (population densities), horizontal transmission produces *βxy* infected individuals. We follow the basic SIR models for modelling horizontal transmission; thus, the transmission of infection is proportional to the densities of the “infected” and that of the “susceptible (non-infected)” individuals (Hethcote, 2000). However, we cannot use the SIR-type models directly since in the present model, the vertical and the horizontal transmission are independent events; the vertical one occurs along a lineage of descendants, while the horizontal one is realized across lineages during the non-reproductive period. Further, in our model, there is no acquired immunity or resistance, i.e., an individual can be cleared anadd then horizontally re-infected several times. For simplicity, we assume that clearing does not depend on individuals’ age, i.e., with *c_r_* ∈ (0,1) an infected host loses its symbiont and becomes non-infected. The uniformity assumptions are crucial and simplifies the model. We can also consider density-dependent competition between infected and non-infected populations for food or other resources, and we assume that both populations are equally effective in the context of this competition. Denote the density dependent competition rate by *γ* ∈ (0,1). Based on these assumptions, we get the following simplified dynamical system,

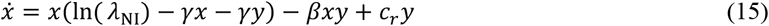

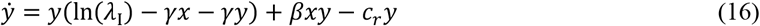

This set of differential equations is implicitly given (Varga et al., 2020) since the Malthusian growth rates ln (*λ*_NI_) and ln (*λ*_I_) are implicitly given by the eigenvalues of the two Leslie matrices of the non-infected and infected host populations, respectively. However, our model is well-defined since there exist unique, positive, and dominant eigenvalues for both Leslie matrices (Caswell, 2001).

Upon analytical investigation, the population size of infected individuals at the positive equilibrium, according to Eq. (16) is,

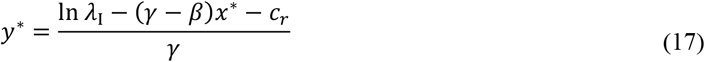

The equilibrium population size *x*\* of the non-infected population is the solution of the following equation,

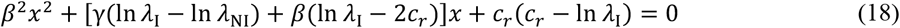

Therefore, the sufficient conditions for the existence of positive fixed points of the resident system (15–16) are ln *λ*_I_ > *c_r_* and *β* > *γ*. Also, this unique interior fixed point is locally asymptotically stable if *γ* < *β* < 3*γ*, and *x*\* and *y*\* are large (see SI (1) for proof). The locally asymptotically stable rest point (*x*\*, *y*\*) of the resident system (15–16) is also globally asymptotically stable (in the sense of Lyapunov, see SI (1) for proof). Figure 4 shows the local dynamics of the resident system around (*x*\*, *y*\*).

**Fig. 4.**
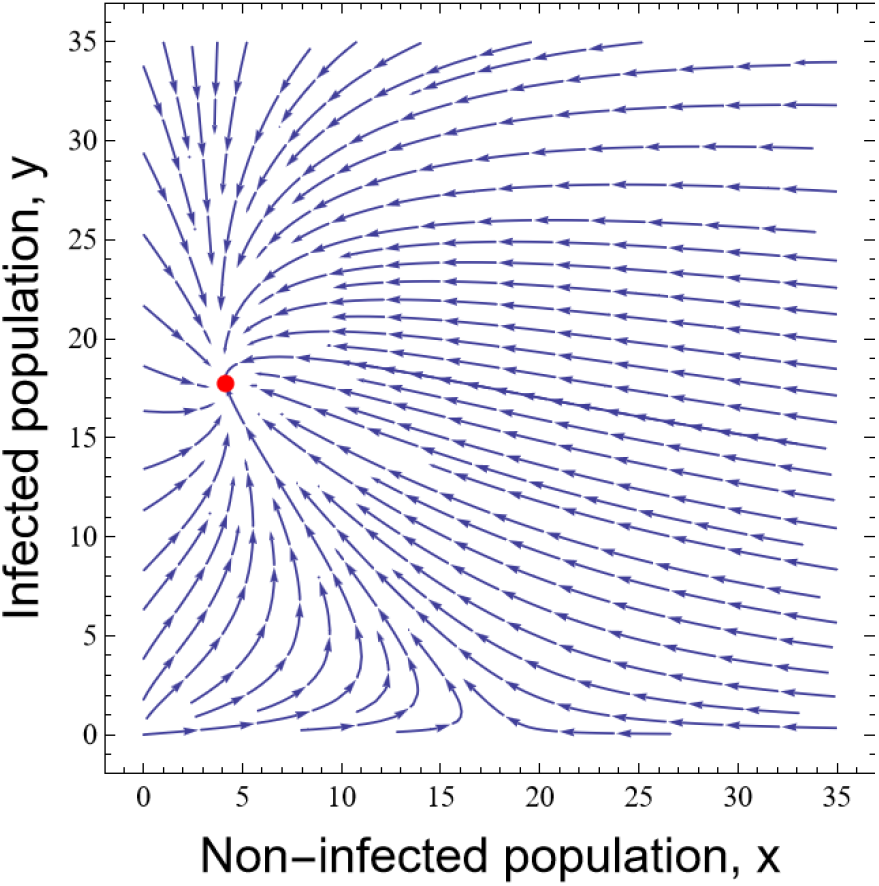
Phase portrait showing the trajectory of the non-infected (*x*) and the infected (*y*) population densities for *λ*_NI_ = 6.303 (from ***L***_NI_(*s*) in Eq. (6) for *s* = 0.544), *λ*_I_ = 9.647 (from ***L***(*s, σ*) in Eq. (14) for *s* = 0.544 and *σ* = 0.492), *β* = 0.15, *γ* = 0.1, and *c_r_* = 0.7. The phase diagram has a stable node at (*x*\*, *y*\*) = (4.131, 17.732)

## 5. Evolutionary stability of the obligate symbiont and the host

We now look for the conditions for the evolutionarily stable ecological state and the uninvadability of rare mutant phenotypes into the resident system. Here we apply the *N*-species evolutionary stability concept of Cressman and Garay 2003a, b. We also implement an often-used simplifying assumption (Ferdy and Godelle 2005; Gandon et al. 2001), i.e., one host has only one symbiont type at a time. In other words, different pathogen types cannot coexist within the same host because different pathogen types displace each other. We are interested in when a rare mutant dies out, and the conditions for evolutionary stability when the possible mutant host is defined by its allocation strategy (*s* ∈ [0,1]) and the possible mutant symbiont is defined by its strategy (*σ* ∈ [0,1]). According to our assumption that the mutation is rare enough, we can consider a single mutant, which appears at the ecologically stable state (*x*\*, *y*\*) of the resident system; thus, a mutation increases the dimension of the system. Note that the stable ecological state (*x*\*, *y*\*) of the resident ecosystem is obtained when the host strategy is *s*\* and the symbiont strategy is σ*. We may as well call the resident equilibrium densities (*x*\*(*s*\*, *σ*\*), *y*\*(*s*\*, *σ*\*)).

Since one of our bio-mathematical aims is to investigate the relationship between the stability of coevolutionary dynamics and game theory (strict Nash equilibrium), we must assume that there is only one mutant at a time in the system according to the definition of Nash equilibrium. Thus, first, we have two cases; 1) when a mutation arises in the symbiont (when *σ* appears), and 2) when a mutation arises in the host (when *s* appears). Finally, we also consider the very rare situation when a mutation arises in both the host and the symbiont species (when both *s* and *σ* appear at the same time). The Malthusian growth rate of the mutant-infected population will change in each case. Based on the condition for the mutant to go extinct, we get the required condition for resisting mutant invasion and evolutionary stability. Figure 5 shows the formation of new mutant systems with the introduction of mutant hosts and symbionts.

**Fig. 5.**
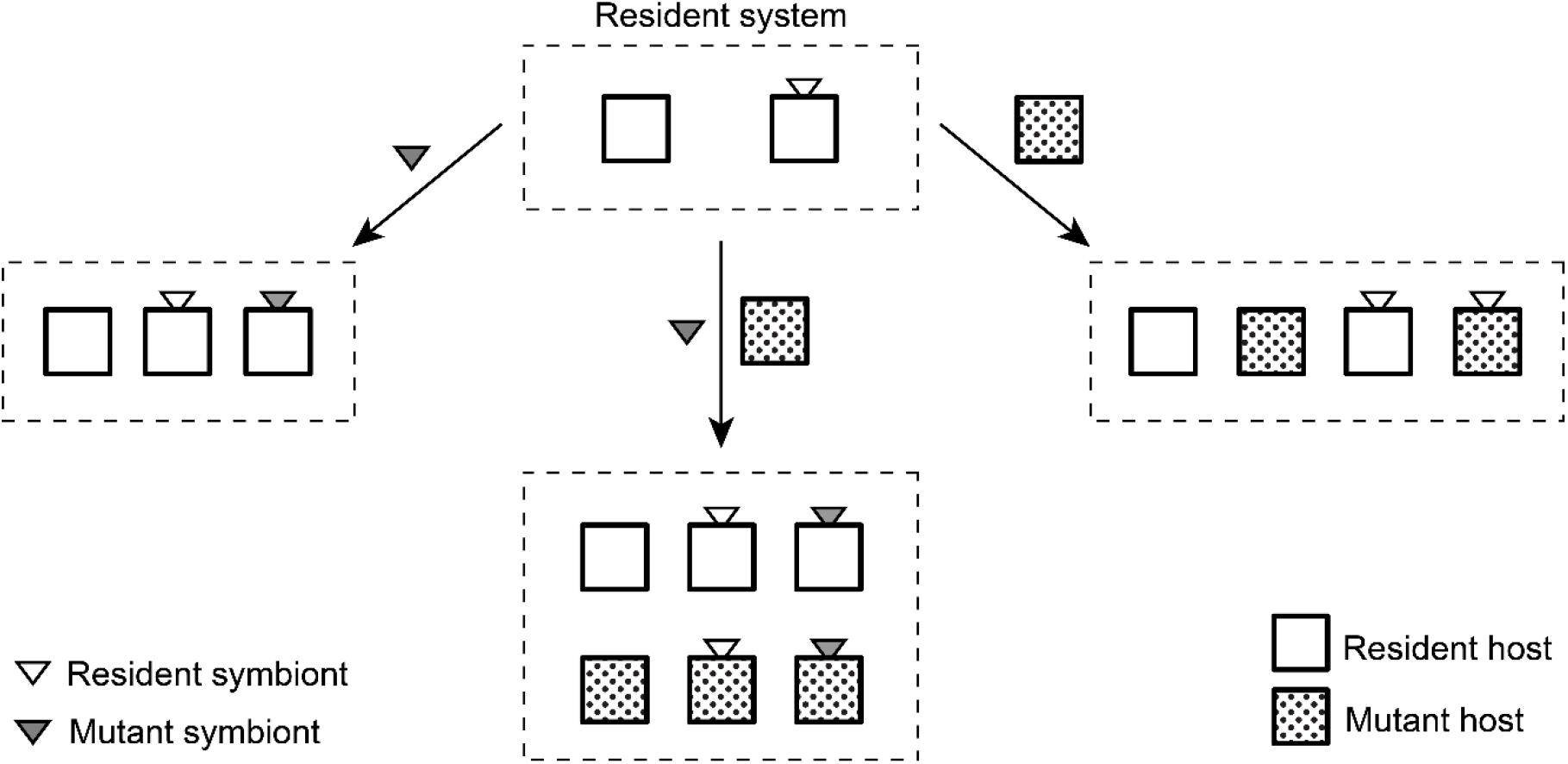
Schematic of mutants introduced into the resident system and the resulting mutant systems with infected and non-infected populations.

### 5.1 Host-symbiont coevolutionary dynamics with mutant symbiont

We first consider the case when a rare mutant arises in the symbiont. For the evolutionary stability of the resident system, we need the mutant to die out by ecological dynamics. Let *z* denote the rare infected mutant population (mutation on the symbiont strategy, *σ* ≠ *σ*\*) with long-term growth rate 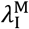. *x* and *y* are as in the resident system. The coevolutionary dynamics of the system with the mutant phenotype of the symbiont is as follows,

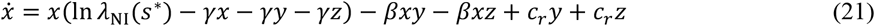

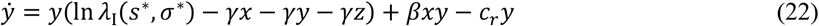

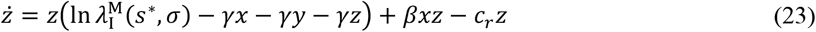

(*x*\*(*s*\*, *σ*\*), *y*\*(*s*\*, *σ*\*), 0) is a positive equilibrium of the system (21) – (23). Sufficient conditions for the given mutant phenotype of the symbiont to die out (mutant cannot invade the stable resident system) are *γ* < *β* < 2*γ* and 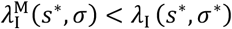 (see SI (2) for proof). In mathematical terms, (*x*\*, *y*\*, 0) is a locally asymptotically stable (l. a. s.) equilibrium of the coevolutionary dynamics (21) – (23) for the above conditions. The condition 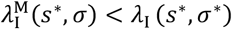 is always true as *λ*_I_ (*s*\*, *σ*\*) is the maximum possible value of *λ*_I_. Figure 6 shows the local dynamics of the system (21) – (23) around (*x*\*, *y*\*, 0).

**Fig. 6.**
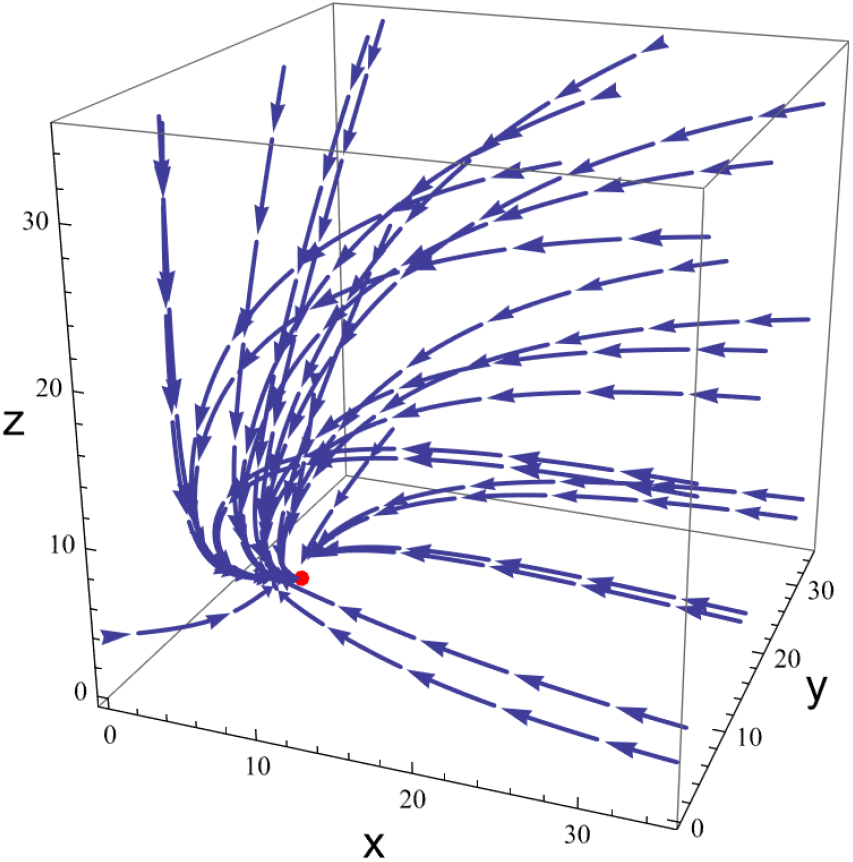
Phase portrait showing the trajectory of the non-infected (*x*), infected (*y*), and mutant-infected (*z*) population densities for *λ*_NI_ = 6.303, *λ*_1_ = 9.647, 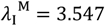, *β* = 0.15, *γ* = 0.1, and *c_r_* = 0.7. The phase diagram has a stable node at (*x*\*, *y*\*, 0) = (4.131, 17.732, 0) that does not allow invasion by the mutant phenotype of the symbiont.

### 5.2 Host-symbiont coevolutionary dynamics with mutant host

Now consider the case when a rare mutant arises in the host. Let *x* denote the density of the non-infected population of resident hosts (strategy *s*\*) and *x*_M_ denote the density of the non-infected population of mutant hosts (strategy *s* ≠ *s*\*). We have only one resident symbiont with strategy*σ*\*, which forms two types of infected populations. Let *y* denote the density of the infected population with the resident host and *z* denote the density of the infected population with the mutant host (strategy *s* ≠ *s*\*). The long-term growth rate of the infected population with mutant host is 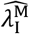. When a non-infected resident (*x*) or mutant (x_M_) encounters any infected populations (either *y* or 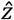), they get infected. The coevolutionary dynamics of the system with the free-living mutant host and infected population with mutant host added to the resident system is as follows,

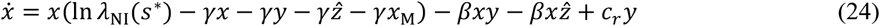

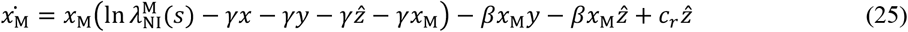

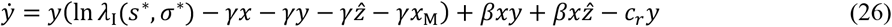

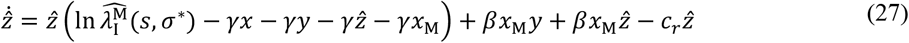

For the evolutionary stability of the resident system, we need the local stability of the positive equilibrium (*x*\*, 0, *y*\*, 0) of (24) – (27). Sufficient conditions for a given mutant phenotype of the host to die out, or in other words, for (*x*\*, 0, *y*\*, 0) to be l. a. s. are 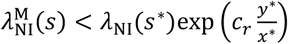 and 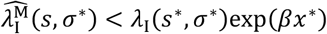 (see SI (3) for proof).

### 5.3 Host-symbiont coevolutionary dynamics with both mutant host and mutant symbiont

Mutations are infrequent; thus, the possibility of independent mutations in the host and in the symbiont strategies at the same time is extremely rare. However, if the densities of hosts and symbionts in the resident system is either extremely high, or not high enough but both species coexist for a very long time, then there is a possibility (extremely rare) for two independent mutations in the hosts and symbionts at the same time. Consequently, in biology the basic assumption of the Nash solution (only one player changes its strategy) does not necessarily hold. The verbal definition of evolutionary stability is as follows. Provided that mutations are rare enough, the resident strategy (*s*\*,*σ*\*) pair is evolutionary stable if coevolutionary dynamics pushes all other possible invading mutants into extinction. The ESS definition is based on the assumption that a single mutation is more probable to occur, i.e., either (*s, σ*\*) or (*s*\*, *σ*) only. However, there is no biological reason to neglect the rare cases when mutations happen in both species at the same time.

Below we consider the case when mutant phenotypes arise in both hosts and symbionts simultaneously. *x, x*_M_, *y, z* and 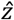 are defined as in system (24) – (27). Let 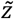 be the infected population with mutant host and mutant symbiont having long-term growth rate 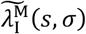. Consequently, we get the following six-dimensional coevolutionary dynamics,

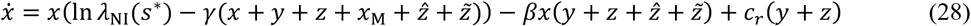

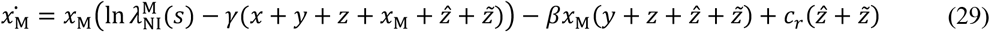

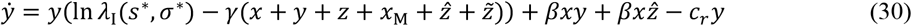

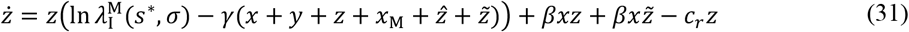

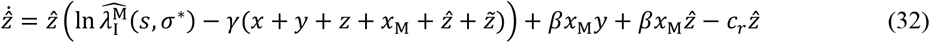

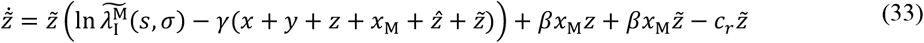

The equilibrium (*x*\*, 0, *y*\*, 0, 0, 0) of the system (28) – (33) with mutations in both host and symbiont is stable if (see SI (4) for proof),

1. 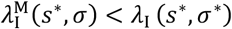,
2. 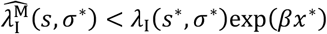
3. 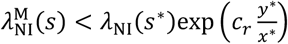, and
4. 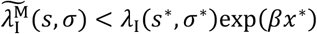

It is worthwhile to note that the ecological aspects of the resident system (*β, c_r_, x*\* and *y*\*) come up in the conditions for evolutionary stability. Conclusively, we observed that the long-term growth rates of the populations govern the conditions for the stability of the resident equilibrium in all the systems. This leads us to question whether our model can be interpreted from the game theory perspective. In the following section, we introduce plausible fitness definitions for both the species (host and symbiont) and try to compare the results of our coevolutionary analysis to that of ESS.

## 6. Intuitive host fitness and Game-theoretic comparison

From the evolutionary game theory perspective, there exists a game-theoretical conflict in our selection situation since the host and the symbiont together determine the infected population’s long-term growth rate. Therefore, we try to compare the Evolutionarily Stable Strategy (ESS) and the linear stability of the coevolutionary dynamics in our model. From a biological point of view, the strict Nash equilibrium (strict Nash implies ESS) means that neither mutant symbionts nor mutant hosts can invade the system (Maynard Smith and Price 1973). For this, we need to define the fitness functions of the players. This problem is intuitively clear for the obligate and vertically transmitted symbionts since their evolutionary success can be given by that of the infected populations. However, the same is not apparent for the hosts. Clearing and horizontal transmission create difficulty since the number of infected and non-infected descendants are dynamically determined in each population. However, we try to provide an intuitive fitness function for the host. Remember that the concept of Nash equilibrium is based on only one player changing its strategy, which means that the mutation (infrequent) occurs only in one of the two species at a time.

We start from the formal definition of the strict Nash equilibrium for an asymmetric game. Consider two players (Host and Symbiont) with strategy sets *s* ∈ [0,1] and *σ* ∈ [0,1] and payoff functions *λ*_H_(*s, σ*) and *λ*_S_(*s, σ*) respectively. Host’s allocation strategy is *s* ∈ [0,1]. Host has a trade-off between reproduction and survival. Symbiont’s strategy is *σ* ∈ [0,1] and it is indicative of its effect on the survival and fecundity of the host.

### Payoff function of Symbiont

Since the symbiont is obligate, its evolutionary success (fitness) is the same as that of the infected population of our model. Thus, its payoff is the long-term growth of the infected population, i.e.,

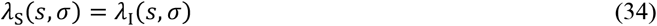

### Intuitive payoff function of Host

Using *λ*_I_ (*s*\*, *σ*\*) and *λ*_NI_(*s*\*) as Malthusian growth rates of each population in the resident system, we can calculate *x*\* and *y*\* (densities at the locally asymptotically stable rest point of the resident dynamics (15–16)). Since the mutation is rare, there is enough time for the dynamics (15–16) to reach its *x*\* stable rest point (*x*\*, *y*\*). Thus, at the resident equilibrium, a host is non-infected or infected with probability 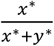 or 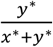, respectively. We define host’s payoff as,

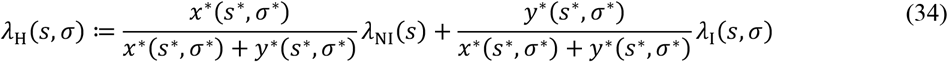

where *λ*_NI_(*s*) and *λ*_I_(*s, σ*) are the long-term growth rates of the non-infected and infected populations respectively, i.e., the dominant eigenvalue of the Leslie matrices ***L***_NI_(*s*) in Eq. (6) and ***L***_I_(*s, σ*) in Eq. (14). Note that the payoffs are not independent. We observe that the host’s payoff function considers both infected and non-infected host lineages. Intuitively, in our model, the long-term growth rate determines the evolutionary success of different phenotypes. According to the mixed transmission mode, each individual host’s lineage will contain infected and non-infected descendants due to clearing and horizontal transmission. Hence, each host individual can alter between infected and non-infected stages throughout its life. Thus, each host’s payoff must consider the long-term growth rates of the infected and the non-infected lineages. The resident system’s rest point determines the ratio of the infected and non-infected descendants at the endpoint of evolution; therefore, we use this ratio to weigh the long-term growth rates of the infected and non-infected lineages. If all hosts are infected at the stable rest point of the resident dynamics, i.e., *x*\* = 0, (say when *c_r_* = 0 and *β* is large enough), then the payoff functions of both players are the same. If most hosts are non-infected (e.g., *c_r_* ≈ 1), then we are near to maximizing *λ*_NI_(*s*).

The host’s best response function *δ*_H_(*σ*) provides the best strategies (which maximize the respective payoff function) that the host must choose when the symbiont plays a given strategy σ. In other words, *δ*_H__H_(*σ*) is a set represented as,

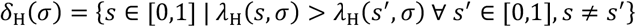

Similarly,

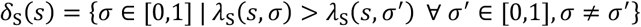

A strict Nash equilibrium is a strategy pair (*s*\*, *σ*\*) such that *s*\* ∈ *δ*_H_(*σ*\* and *σ*\* ∈ *δ_S_*(*s*\*). where *s*\* and *σ*\* are the best strategies of the host and the symbiont respectively (see Figure 7). Equivalently in our model, a strategy pair (*s*\*, *σ*\*) is an ESS (strict Nash equilibrium) if for all *s* ≠ *s*\* and *σ* ≠*σ*\*, we have 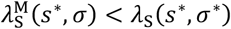 and 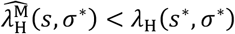, we could arrive at the following inequalities,

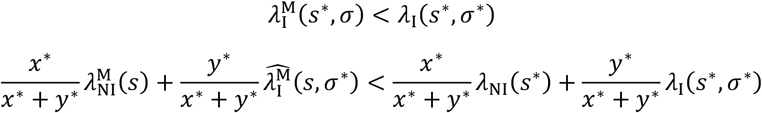

**Fig. 7.**
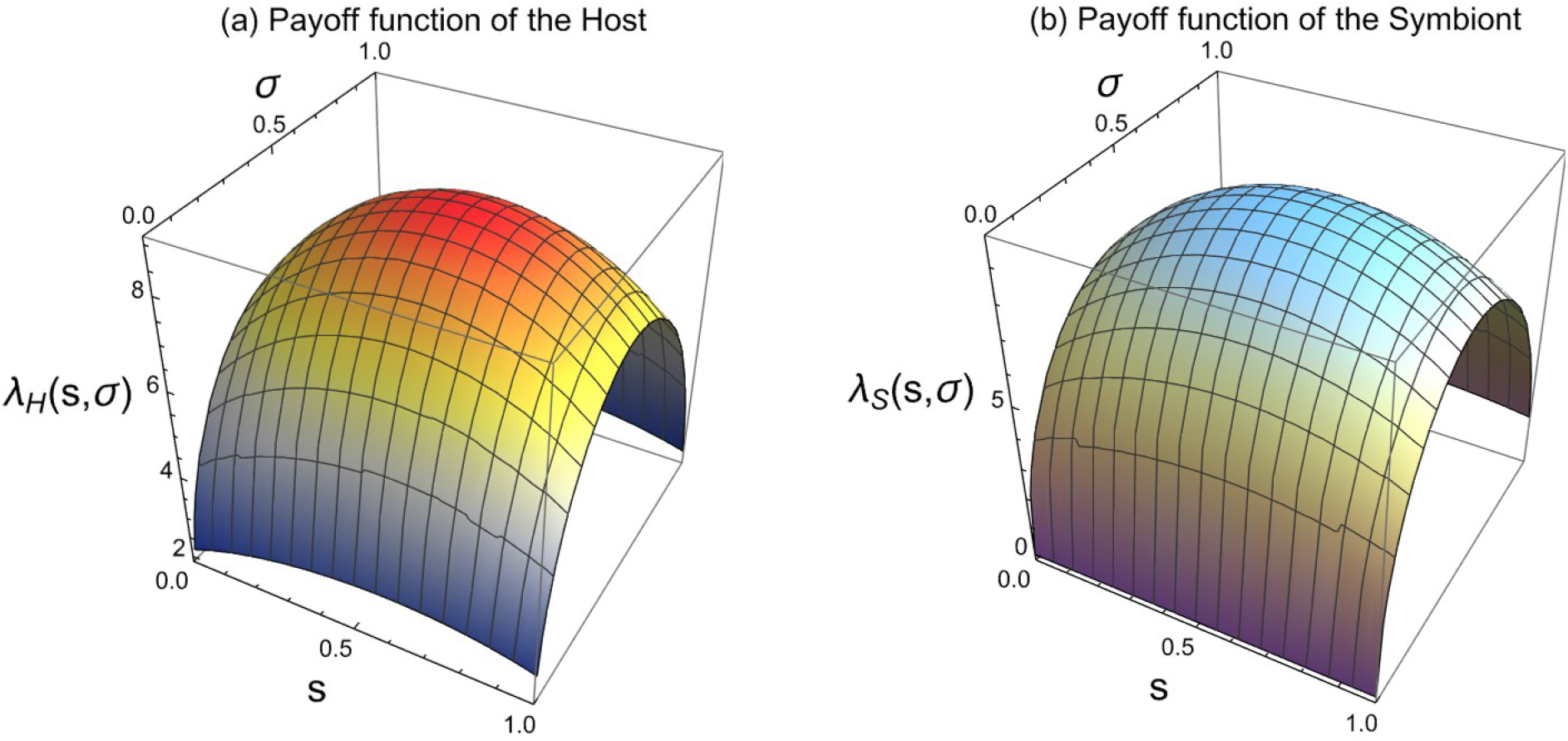
The payoff function of the host and the symbiont, respectively, on *s* – *σ* plane. Observe that the payoff functions *λ*_H_(*s, σ*) and *λ*_S_(*s, σ*) are continuous and strictly concave-down functions with a maximum at a single point. Analyzing the plots, the best response function of host *δ*_H_(*σ*) maps to a singleton set with element *s*\* = 0.544 and the best response function of symbiont *δ_s_*(*s*) maps to a singleton set with element *σ*\* = 0.492. Therefore, the strict Nash equilibrium is (*s*\*, *σ*\*) = (0.544, 0.492).

Thus, in our model, game theoretic analysis based on the intuitive host fitness function gives coarser conditions compared to the following conditions for stability from our coevolutionary dynamical model,

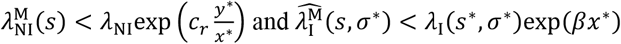

Note that the definition of strict Nash equilibrium of the above game is strictly based on our uniform simplifying assumptions (i.e., clearing, horizontal transmission, and competition between the infected and non-infected resident/mutant populations are independent of the strategies), and there is only one mutant at a time. To define an evolutionary game, we must give the average fitness of different phenotypes belonging to different species. However, that is not obvious in our above model for the hosts since each individual can switch from infected to non-infected and vice-versa during its life. Our main observation is that ecological coevolutionary dynamics requires more stringent conditions than the game-theoretical method. However, this conjecture needs more studies in more general selection situations when our uniformity conditions are not satisfied.

## 7. Discussion, interpretation, and biological examples

In this manuscript, we introduced a combination of the Leslie demography model (vertical transmission) and competitive selection dynamics (horizontal transmission with clearing) to explore the long-term growth rate of symbiont-infected hosts against non-infected hosts. An unusual feature of the model is that the appearance of two mutants (mutant host and mutant symbiont) in the coevolutionary system increased its dimensionality to six. Then we applied a game-theoretical approach to the same model as an alternative method. From a purely methodological point of view, we observed that the game-theoretical approach yields coarser results than the analysis of coevolutionary dynamics for two reasons. First, the strict Nash definition allows only one mutant phenotype at a time, either in the host or in the symbiont. Second, coevolutionary dynamics performs better even if it is limited to one mutant type as. we were able to obtain more stringent and precise conditions for the uninvadability of mutants. From a biological point of view, our model yields both trivial and non-trivial connotations. Evolutionary theory predicts that uniparentally and vertically transmitted parasites must be harmless to their hosts (Fine 1975; Yamamura 1993; Lipsitch et al. 1995). Thus, obligate symbiosis with vertical transmission will likely end up as mutualism. Accordingly, a symbiont species may not develop or maintain a parasitic way of life (i.e., may not decrease host fitness) in a uniparental system unless capable of horizontal transmission (Garay et al. 2016). This means parasites that utilize only the parent-offspring (vertical) transmission routes may not survive because the non-infected lineages outcompete the infected ones since it has a better long-term growth rate. Presuming that the bodily contacts between parents and offsprings are usually more direct and more long-lasting than between other conspecifics - thus more proper for symbiont transmission - this means that mutualism is supposed to be an archaic form of symbiosis. Parasitism is likely to be a derived way of life that can emerge only after the symbiont has already evolved the more advanced ways of horizontal transmission. Alternatively, the parasitic way of life may evolve from a sapronotic life strategy (Kuris et al. 2014).

We observed that the evolutionary success of an obligate symbiont and its host depends on at least three factors; 1) the symbiont’s effect on the host’s life history, 2) the rate of clearing of infection, and 3) the frequency and mode of transmission. The connection between obligate symbiont*s*\* effects on host*s*\* life history and their transmission mode is well explored in both theoretical and empirical studies (Bibian et al. 2016; Brown and Akçay 2019; Chung et al. 2015; Clayton et al. 2015; Ebert 2013; Ewald 1987; Gandon et al. 2008; Rudgers et al. 2012). In the standard models, these effects are on host survival and fecundity (Ferdy and Godelle 2005). Further, the trade-offs between host survival and fecundity have been shown theoretically to determine the persistence of symbioses (Bibian et al. 2016; Chung et al. 2015; Rudgers et al. 2012; Yule et al. 2013). Note that the symbiont*s*\* effects on the host*s*\* life history can be more complex, particularly in hosts having a long and complex life cycle with different developmental stages.

During the coevolutionary process, not only the symbiont strategy but also the host strategy can modify the infected host*s*\* life-history parameters. To get an insight, we strictly focused on how the two partner*s*\* strategies together modify the infected population’s life-history traits. For this purpose, we applied simplifying assumptions. Hence, our conceptual model was strictly based on uniformity conditions (i.e., clearing, transmission mode, and competitive ability are independent of host and symbiont strategies). These assumptions ensure the following advantages:

1. They radically decrease the dimension of the dynamical system (Hancock et al., 2011) or the integral projection model’s dimension (Chung et al., 2015; Yule et al., 2013). For instance, if the host has three developmental stages (say with different horizontal transmission rates), we get a six-dimensional resident system. Further, if obligate mutant symbionts and mutant hosts are introduced, we have nine-and twelve-dimensional coevolutionary dynamics, respectively.
2. The main novelty of our present conceptual model is that it gives an insight into cases when the host has several developmental stages, and the symbiont can manipulate the survival and fecundity of the different stages in different ways.
3. They separate the density-dependent phenomena (like competition and horizontal transmission) from the host-symbiont interaction within the host body.

Our model also provides a close insight into the complexity of the concept of virulence. Classical textbook wisdom says that virulence is the infection-induced reduction of host survival and/or reproductive success (Anderson and May 1978). However, our model shows that symbiont*s*\* effect on host longevity and reproduction may be different, even opposite, and their net effects may often be counterintuitive. To exemplify the potential complexity of host-symbiont interactions, below we list eight categories based on the infection*s*\* effects on host longevity and reproduction success, with biological examples:

1. **Survival-reducers:** the Japanese subgroup of Human T-cell lymphotropic virus type 1 (HTLV-I) can serve as an example of this category. It is often transmitted from mother to child through breastfeeding and is frequently associated with a highly lethal disease, the adult T-cell leukaemia. However, since the onset of the disease is about sixty years (Murphy et al. 1989), we can reasonably assume that it reduces only host longevity without affecting its reproductive success.
2. **Fecundity-reducers:** cytoplasmic incompatibility induced by Wolbachia infections in insects (Sinkins 2004) nicely exemplify this category. Note that reduced host fecundity may either result from pathogenic effects or, alternatively, as a damage limitation strategy of the host (Hurd 2001).
3. **Survival-and-fecundity-reducers:** several well-known parasites and pathogens reduce host longevity and reproductive success in parallel.
4. **Survival-increasers:** imagine a hypothetical symbiont that increases host longevity without affecting lifetime reproductive success. We could not find examples of such organisms in the literature.
5. **Fecundity-increasers:** Wolbachia infections which increase male fertility in the beetle in *Tribolium confusum* (Wade and Chang 1995). Another intracellular parasitic bacterium, the so-called Cytophaga-Like Organism (CLO), increases female fertility without influencing the mortality of the infected host, the predatory mite *Metaseiulus occidentalis* (Weeks and Stouthamer 2004).
6. **Survival-and-fecundity-increasers:** Several well-known mutualists belong to this category.
7. **Survival-increaser-and-fecundity-reducer:** The larvae of several helminths (like Cestodes, Trematodes, and Acanthocephalans) destruct host gonads (a phenomenon called ‘parasitic castration’) apparently to increase host body size (‘parasitic gigantism’) and also survival (Berec and Maxin 2012). Fungal endophytes decrease fecundity and increase survival for the grass Poa alsodes (Chung et al. 2015). Larvae of the rat tapeworm *Hymenolepis diminuta* also increases the survival of the intermediate host (the mealworm beetle *Tenebrio molitor*) at the expense of reducing its fecundity (Hurd et al. 2001).
8. **Survival-reducer-and-fecundity-increaser:** fungal endophytes increase fecundity in the grass *Agrostis hyemalis* at the expense of reducing its survival (Yule et al. 2013). *Toxoplasma gondii*, a widespread human infection, may cause flu-like illness and various neuropsychiatric symptoms. On the other hand, infections make patients sexually more attractive. Since they have more sexual partners, their reproductive success may increase (Borráz-León et al. 2022)

Evidently, the idea that symbionts may affect host lifespan and fecundity differently is not new (Brown and Akçay 2019). Therefore, we aimed to develop a model that enables us to handle the complexity of host-symbiont interactions. For this purpose, we have generalized our former Kin Demographic Selection Model (Garay et al. 2016; Garay et al. 2018; Garay et al. 2018) where we only considered mutant and resident lineages that competed with each other. Contrarily, here we have focused on two interacting species with uniform interaction patterns and horizontal infection routes between infected and non-infected lineages. The novelty of this general model is that horizontal infection connects the resident and mutant infected lineages. This opens future avenues to build more complex models with other features like evolving or strategy-dependent horizontal transmission or clearing or models with symbionts that increase host competitive abilities. In this sense, we incorporate different aspects of ecology to study resistance to mutant invasion and evolutionary stability.

## Acknowledgements

N. K. and J. G. acknowledge funding received from the European Union’s Horizon 2020 Research and Innovation Programme under the Marie Skłodowska-Curie grant agreement number 955708. L. R.’s work was supported by the National Research, Development and Innovation Office in Hungary (RRF-2.3.1-21-2022-00006) and the National Research, Development and Innovation Fund of Hungary (K143622).

## Author contributions

N. K.: Methodology, Formal analysis, Visualization and Writing – Original draft; L. R.: Conceptualization; A. S.: Methodology; J. G.: Conceptualization, Methodology and Supervision; all authors read and approved the manuscript.

## Statements and Declarations

### Competing interests

The author(s) declare no competing interests.

## SUPPLEMENTARY INFORMATION

### SI (1) Stability of the Resident System

We first conduct the stability analysis of the equilibrium points corresponding to the following set of differential equations (15) – (16) in the main text,

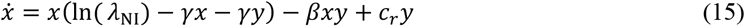

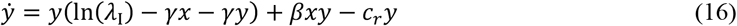

The equilibrium population size of the infected population,

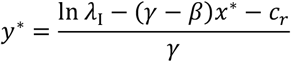

The equilibrium population size *x*\*of the non-infected population is the solution of the following equation,

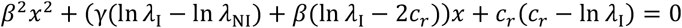

The roots are positive real if ln *λ*_I_ > *c_r_*, i.e., the Malthusian growth rate of the infected population is larger than the clearing rate. This relation also guarantees that there is at least one positive *x*\*. Also, *y*\* is positive if *β* > *γ*. Therefore, the sufficient conditions for the existence of positive rest points of (15) – (16) are ln *λ*_I_ > *c_r_* and *β* > *γ*.

#### Theorem 1

*The rest point* (*x*\*, *y*\*) *of the resident system (15)* – *(16) is locally asymptotically stable for ln λ_I_* > *c_r_, γ* < *β* < 3*γ, large x*\* *and large y*\*.

*Proof:* Consider the fixed point of the above dynamics in the positive quadrant, i.e., 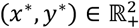 The Jacobian matrix at this fixed point is,

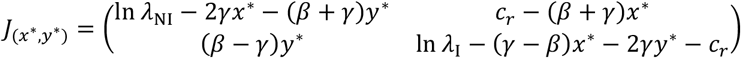

The local stability of the rest point (*x*\*, *y*\*) is implied by the following two inequalities: tr *J*_(*x*\*, *y*\*)_ < 0 and det *J*_(*x*\*, *y*\*)_ > 0. The trace is negative if ln *λ*_NI_ + ln *λ*_I_ – *c_r_* – (3*γ* – *β*)*x*\* – (3*γ* + *β*)*y*\* < 0. Large equilibrium densities *x*\*, *y*\* and *β* < 3*γ* make the negativity possible. The large *x*\* and *y*\* also help to satisfy the positivity condition of the determinant by making *J*_11_, *J*_12_ and *J*_22_ negative, while *J21* is positive because of *β* > *γ*. This implies a 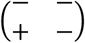 sign pattern resulting in a positive determinant. To sum up, the following conditions make the existence and locally stability of the positive equilibrium (*x*\*, *y*\*) possible,

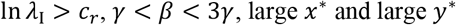

In other words, (*x*\*, *y*\*) is a stable (locally asymptotically) node for the above conditions.

#### Global stability of the Resident system

##### Theorem 2

*The locally asymptotically stable rest point* (*x*\*, *y*\*) *of the resident system (15)* – *(16) is also globally asymptotically stable*.

*Proof:*

Consider the resident system,

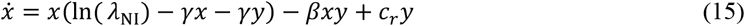

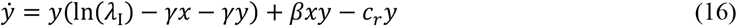

Let (*x*\*, *y*\*) be the unique interior equilibrium of the dynamical system. The conditions for the existence of the locally asymptotically stable unique interior equilibrium (as mentioned in the previous section) of the resident system are,

1. ln *λ*_I_ > *c_r_*,
2. *γ* < *β* < 3*γ*, and
3. large *x*\* and large *y*\*

We now use the Lyapunov direct method to conduct global stability analysis. Consider the following Lyapunov function,

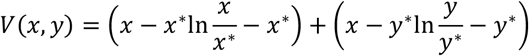

The candidate Lyapunov function is radially unbounded as the function is increasing for *x* > *x*\*, *y* > *y*\*.

Also, *V*(*x*\*, *y*\*) = 0 and *V*(*x, y*) > 0 for (*x, y*) ≠ (*x*\*, *y*\*). Time derivative of the Lyapunov function is,

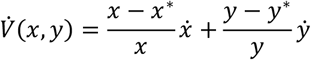

For asymptotic stability, we need to prove that 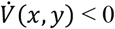. Let 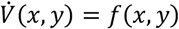, hence *f*(*x*\*, *y*\*) = 0.

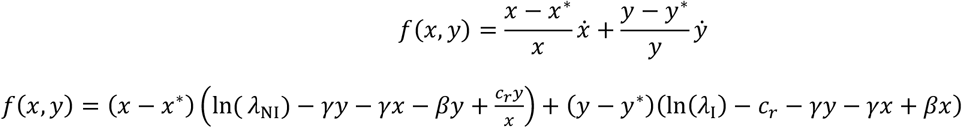

Differentiating *f*(*x, y*) w.r.t *x* and *y*,

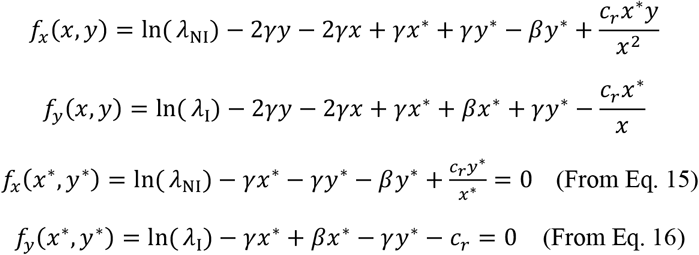

(*x*\*, *y*\*) is the unique interior equilibrium point of the dynamics (15) – (16) and is also the unique critical point of *f*(*x, y*) for *x* and *y* > 0. The Hessian matrix corresponding to *f*(*x, y*) at (*x*\*, *y*\*) is,

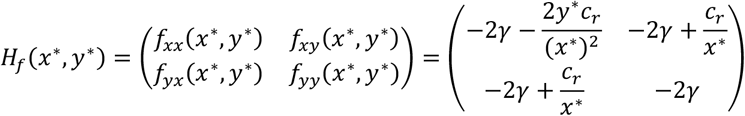

We see that *H_f_*(*x*\*, *y*\*) is negative definite (all eigenvalues are negative) if *c_r_* < 4*γ*(*x*\* + *y*\*). Since *x*\*, *y*\* are large this is possible. Therefore, the unique critical point (*x*\*, *y*\*) is a maximum point of *f*(*x, y*).

Thus, for any (*x, y*) ≠ (*x*\*, *y*\*), *f*(*x, y*) < *f*(*x*\*, *y*\*). Since *f*(*x*\*, *y*\*) = 0, we can conclude,

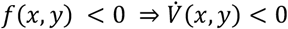

Therefore, by Lyapunov stability criterion, (*x*\*, *y*\*) is globally asymptotically stable.

**Fig. SI 1.1.**
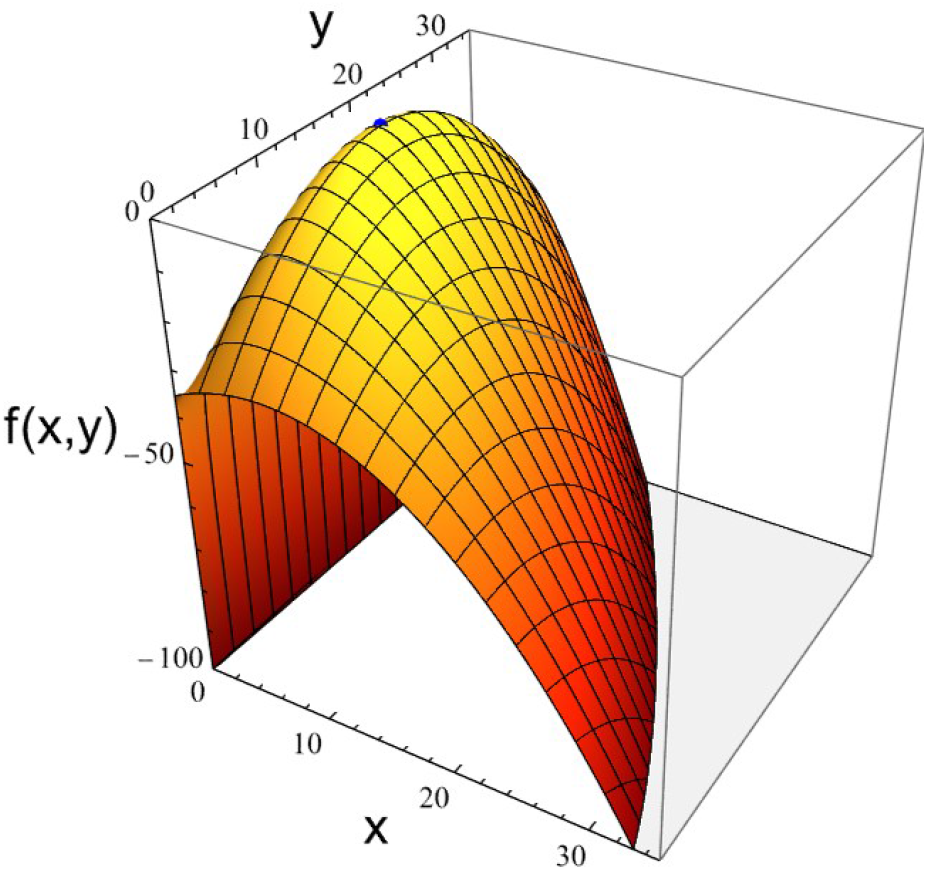
The time derivative of the Lyapunov function, 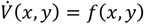 attains maximum value at (*x*\*, *y*\*) = (4.131, 17.732) (blue point) for *λ*_NI_ = 6.303, *λ*_I_ = 9.647, *β* = 0.15, *γ* = 0.1, and *c_r_* = 0.7. Since 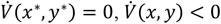 for all (*x, y*) ≠ (*x*\*,*y*\*)

### SI (2) Stability analysis of host-symbiont coevolutionary dynamics with mutant symbiont

Consider the following coevolutionary dynamics of the system (21) – (23) with the mutant phenotype of the symbiont, where *z* denote the infected population with mutant symbiont (strategy *σ* ≠ *σ*\*) with long-term growth rate 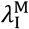, and *x* and *y* are as in the resident system,

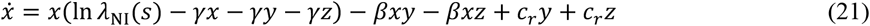

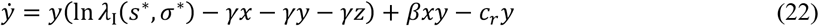

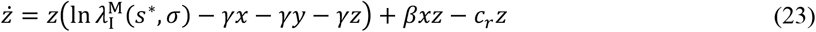

#### Theorem 3

*If for a given rare infected mutant population with mutant symbiont, we have 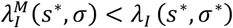, then this mutant phenotype cannot invade the resident system*.

*Proof:* For the condition when mutants die out in this case, i.e., (*x*\*, *y*\*, 0) is locally asymptotically stable, we investigate the Jacobian matrix of the dynamics (21) – (23) at (*x*\*, *y*\*, 0), which reads as,

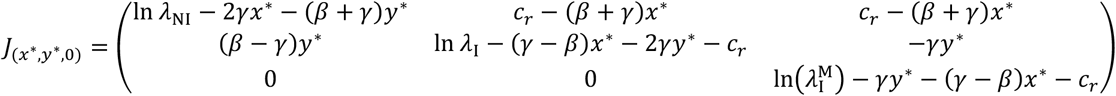

Recall from the stability analysis of the resident system (16) – (17), we have ln *λ*_I_ > *c_r_* and 3*γ* > *β* > *γ*, for the existence of (*x*\*, *y*\*).

According to Routh Hurwitz criterion (the necessary and sufficient conditions for the linear stability of the system), the conditions for the local asymptotic stability of the equilibrium point are,

1. tr *J*_(*x*\*, *y*\*, 0)_ < 0,
2. det *J*_(*x*\*, *y*\*, 0)_ < 0,
3. 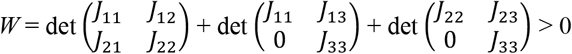
4. tr *J*_(*x*\*, *y*\*, 0)_ * *W* < det *J*_(*x*\*, *y*\*, 0)_

The trace is negative if 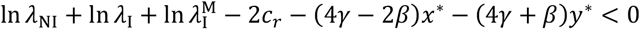. Large equilibrium densities (*x*\*, *y*\*) and *β* < 2*γ* make the negativity possible. The conditions ln *λ*_I_ > *c_r_*, *γ* < *β* < 3*γ*, large *x*\* and large *y*\* make the det 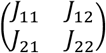 positive as stated in the stability analysis of the resident system. Therefore, det *J*_(*x*\*, *y*\*, 0)_ < 0 if *J*_33_ < 0, i.e.,

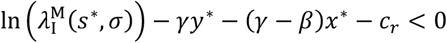

Also, 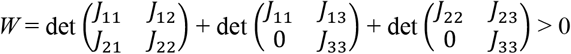 if *J*_33_ < 0.

tr *J*_(*x*\*, *y*\*, 0)_ < 0, det *J*_(*x*\*, *y*\*, 0)_ < 0, *W* > 0 and tr *J*_(*x*\*, *y*\*, 0)_ * *W* < det *J*_(*x*\*, *y*\*, 0)_ for *γ* < *β* < 2*γ*, large *x*\*, large *y*\* and ln 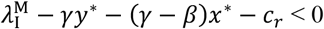.

Sufficient conditions for a given mutant symbiont to die out are large equilibrium densities (*x*\*, *y*\*), *γ* < *β* < 2*γ*, ln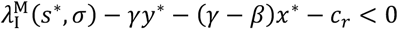 (or equivalently 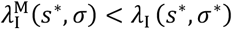, see below).

Reinterpreting the stability condition,

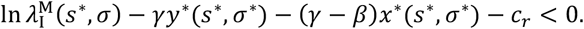

where 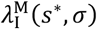 is the dominant eigenvalue of the Leslie matrix corresponding to the infected lineage at (*s*\*, *σ*). Moreover (*x*\*(*s*\*, *σ*\*), *y*\*(*s*\*, *σ*\*)) is the rest point of the resident two-dimensional ecological dynamics (15) – (16).

From the rest point of the resident ecosystem (15) – (16), we have,

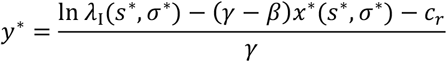

Thus, for the evolutionary stability of the strategy pair (*s*\*, *σ*\*) for all strategy pair (*s, σ*) ≠ (*s*\*, *σ*\*) we need,

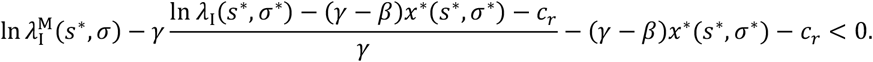

After some elementary algebra we get,

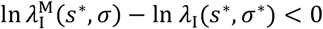

Since the function ln is strictly monotone, (*s*\*,*σ*\*) is evolutionary stable, if for other strategy pair (*s, σ*) ≠ (*s*\*, *σ*\*) we have,

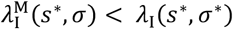

### SI (3) Stability analysis of host-symbiont coevolutionary dynamics with mutant host

Consider the coevolutionary dynamics with infected population having mutant hosts. The coevolutionary dynamics of this four-dimensional system is as follows,

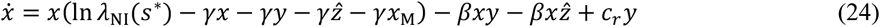

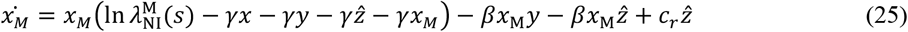

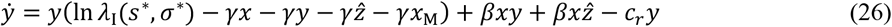

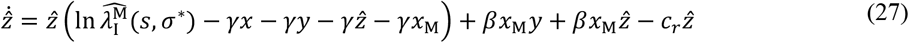

#### Theorem 4

*If for a given rare infected mutant population with mutant host, 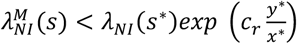 and 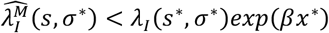, then this mutant cannot invade the stable resident system. In mathematical terms*, (*x*\*, 0, *y*\*, 0) *is a l.a.s. fixed point of the coevolutionary dynamics (24)-(27)*.

*Proof:* We investigate the Jacobian matrix of dynamics (24) – (27) at (*x*\*, 0, *y*\*, 0), which reads as,

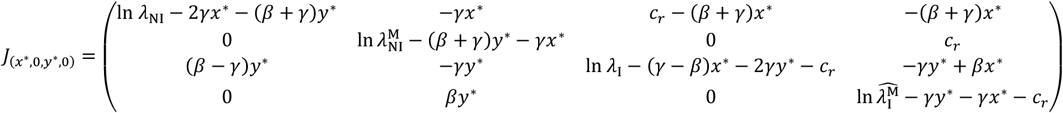

The eigenvalues of *J*_(*x*\*, 0, *y*\*, 0)_ are:

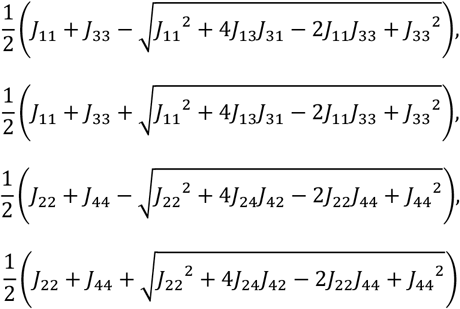

All the eigenvalues are negative if *J*_11_, *J*_22_, *J*_33_, *J*_44_ are negative and the following inequalities hold *J*_13_*J*_31_ < *J*_11_*J*_33_, and *J*_24_*J*_42_ < *J*_22_*J*_44_. From the stability of the resident system (15) – (16) and comparing the elements of the Jacobian matrices, we can observe that *J*_11_, *J*_33_ < 0 and *J*_13_*J*_31_ < *J*_11_*J*_33_. Also, for sufficiently large *x*\*, *y*\*, as in the resident system, *J*_24_*J*_42_ < *J*_22_*J*_44_.

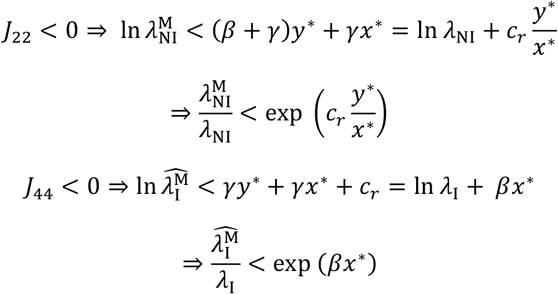

The rest point (*x*\*, 0, *y*\*, 0) is locally asymptotically stable if 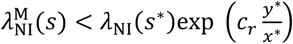 and 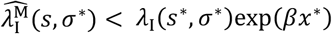.

### SI (4) Stability analysis of host-symbiont coevolutionary dynamics with mutant host and mutant symbiont

Consider the following six-dimensional coevolutionary dynamics for the case when mutant phenotypes are introduced in both the host and the symbiont populations,

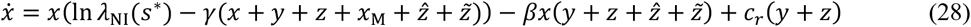

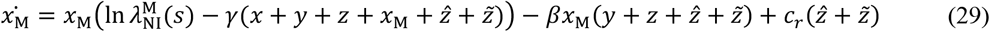

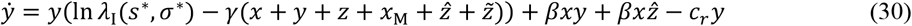

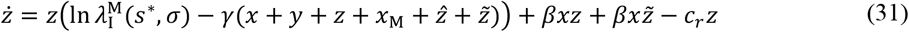

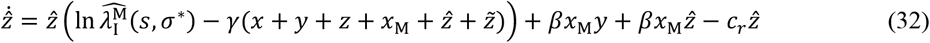

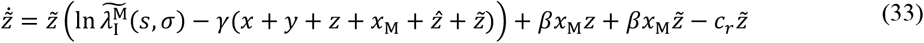

#### Theorem 5

*The rest point* (*x*\*, 0, *y*\*, 0, 0, 0) *of the system (28)* – *(33) with both mutant hosts and mutant symbionts is stable if 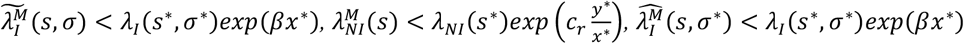, and 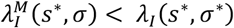*.

*Proof:* We investigate the Jacobian matrix of dynamics (28) – (33) at (*x*\*, 0, *y*\*, 0, 0),

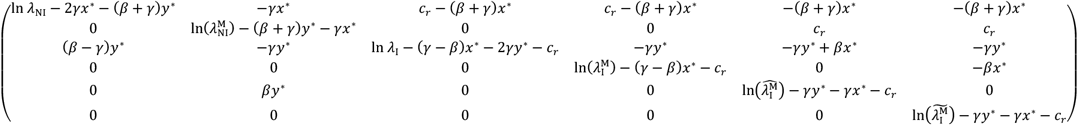

The corresponding characteristic polynomial is (if *μ* is an eigenvalue),

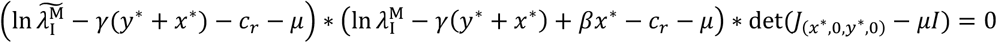

The system with both mutant hosts and mutant symbionts is stable at (*x\**, 0, *y\**, 0, 0) for the following conditions,

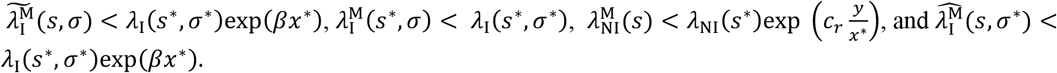

## Notes

### Competing Interest Statement

The authors have declared no competing interest.

